# EEG evidence that morally relevant autobiographical memories can be suppressed

**DOI:** 10.1101/2022.03.01.482486

**Authors:** Akul Satish, Robin Hellerstedt, Michael C. Anderson, Zara M. Bergström

## Abstract

Remembering unpleasant events can trigger negative feelings. Fortunately, research indicates that unwanted memories can be suppressed to prevent them from intruding into awareness, improving our mental state. The current scientific understanding of memory suppression is, however, based mostly on simpler memories such as associations between words or pictures, which may not reflect how people control unpleasant memory intrusions in everyday life. Here, we investigated the neural and behavioural dynamics of suppressing personal and emotional autobiographical memories using a modified version of the Think/No-Think task. We asked participants to suppress memories of their own past immoral actions, which were hypothesised to be both highly intrusive and motivating to suppress. We report novel evidence from behavioural, ERP and EEG oscillation measures that autobiographical memory retrieval can be suppressed and suggest that autobiographical suppression recruits similar neurocognitive mechanisms as suppression of simple laboratory associations. Suppression did fail sometimes, and EEG oscillations indicated that such memory intrusions occurred from lapses in sustained control. Importantly however, participants improved at limiting intrusions with repeated practice. Furthermore, both behavioural and EEG evidence indicated that intentional suppression may be more difficult for memories of our morally wrong actions than memories of our morally right actions. The findings elucidate the neurocognitive correlates of autobiographical retrieval suppression and have implications for theories of morally motivated memory control.

---

When faced with reminders of disturbing past events, memories of the event and associated negative emotions can intrude into awareness and lead to an unpleasant state of being. Fortunately, people can recruit cognitive control processes to suppress retrieval of unwanted memories and consequently feel better (Anderson & Hanslmayr, 2014). This ability to exert memory control is an important aspect of maintaining good mental health (Engen & Anderson, 2018; Mary et al., 2020), and a large body of research over the last 20 years has delineated, with increasing detail, the neurocognitive mechanisms that enable memory suppression. However, most research studies the suppression of memories of simple stimuli, such as words or pictures encoded in a laboratory environment (Anderson & Hanslmayr, 2014). Research on the neural basis of memory control of real-world, emotionally charged autobiographical memories (Noreen et al., 2016) remains limited, despite the theoretical view that memory suppression is a motivated process that people use to regulate emotions. Here, we used EEG to investigate the neurocognitive mechanisms involved in suppressing autobiographical memories of events that people should be particularly motivated to avoid – specifically, memories of behaving in a morally wrong manner (Stanley & De Brigard, 2019).

Memory control is typically studied in the laboratory using the Think/No-Think (TNT) paradigm (Anderson et al., 2004; Anderson & Green, 2001). Participants initially learn pairs of stimuli, such as two weakly related words (e.g., ordeal-roach). Next, one stimulus in each pair is presented as a reminder for the other. Participants are instructed to actively remember and keep in mind the associated stimulus if the reminder is displayed in green (Think items), but if it is shown in red (No-Think items), they are instructed to prevent the memory of the associated stimulus from intruding into consciousness. The same suppression or retrieval task is repeated many times for each reminder. Participant ratings of whether or not the memory came to mind for each Think/No-Think trial (Levy & Anderson, 2012) are used to assess how often No-Think memories intrude into conscious awareness despite participants’ attempts at stopping retrieval. A reliable finding is that such intrusions are less frequent over repeated suppressions (Benoit et al., 2015; Davidson et al., 2020; Gagnepain et al., 2017; Harrington et al., 2021; Hellerstedt et al., 2016; Levy & Anderson, 2012; Mary et al., 2020; van Schie & Anderson, 2017), providing real-time evidence that retrieval suppression is increasingly effective the more times it is applied to a memory. Following the TNT task, participants complete a surprise memory test of all cue-target pairs, wherein the reminders are presented again and participants are asked to recall and report all associated words (or pictures), regardless of previous TNT instructions. Memories in the No-Think condition are typically more poorly recalled on the final test compared to both memories in the Think condition and memories in a Baseline condition that were neither repeatedly suppressed nor retrieved, suggesting that intentional suppression can induce subsequent forgetting (Anderson & Hanslmayr, 2014; Stramaccia et al., 2020).

EEG measurements during the Think/No-Think phase have been used to elucidate the neurocognitive mechanisms involved in memory suppression with high temporal resolution. Previous ERP findings indicate that attempting to suppress retrieval in response to a reminder involve early attentional and cognitive control processes between around 200-500ms (Bergström et al., 2009b; Crespo-García et al., 2021; Mecklinger et al., 2009; Streb et al., 2016; Waldhauser et al., 2012). When suppression is successful, cognitive control reduces the ERP marker of conscious recollection, which is evidenced by a reduced parietal positivity for supressed (No-Think) memories compared to retrieved (Think) memories that is maximal between 500-800ms (Bergström et al., 2007, 2009a, 2009b; Depue et al., 2013). From that time onwards, sustained control needs to be maintained to ensure that the memory does not enter into awareness for as long as the person is exposed to the reminder (Hanslmayr et al., 2009). Previous EEG oscillation studies indicate that sustained memory control is associated with a decrease in oscillatory power for suppressed compared to retrieved memories across theta (~4-7Hz), alpha (~8-12Hz), and beta (~13-30Hz) frequency bands. It has been suggested that parietal theta power reductions reflect reduced recollection of the associated memory^1^, whereas sustained maintenance of cognitive control to ensure that the memory does not enter awareness is reflected by alpha/beta band reductions (Depue et al., 2013; Legrand et al., 2020; Quaedflieg et al., 2020; Waldhauser et al., 2015).

One prior study has investigated the ERP correlates of self-reported intrusions (Hellerstedt et al., 2016). In that study, intrusions were associated with an early fronto-central ERP positivity around 400ms after the cue, potentially indicating initial reactivation of a memory trace (Hellerstedt et al., 2016), followed by a negative-slow-wave that was maximal between 550-900ms. The negative slow-wave was suggested to reflect the intruding memories being active in working memory, or alternatively, processes related to error detection. Contrary to predictions, the study did not show enhanced parietal positive ERP markers of conscious recollection during intrusions. Measuring EEG oscillatory correlates of memory intrusions, (Castiglione et al., 2019) found that successful stopping of both memory retrieval and motor actions was associated with early (200-300ms) increased right-frontal beta oscillation power (~13-30Hz), which decreased when participants experienced memory intrusions. These results therefore suggested that intrusions occurred when participants failed to engage inhibitory control mechanisms that were reflected by beta band oscillations.

Prior research, however, has mostly used simple stimuli, such as pairs of words or images to investigate the neurocognitive basis of memory suppression. Although such stimuli are useful for achieving experimental control and for understanding the core processes involved in the task, they do not address how people control retrieval of more complex, self-relevant, and emotional memories in everyday life. There is extant behavioural and neuroimaging literature on suppression of emotional words/picture memories (e.g., Chen et al., 2012; Depue et al., 2007; Gagnepain et al., 2017; Joormann et al., 2009; Legrand et al., 2020), but because of the lack of self-relevance of those materials, they arguably do not elicit strong motivation for memory control. Research using more complex stimuli suggests that people can suppress unpleasant self-relevant information (Benoit et al., 2016; Noreen et al., 2014) and autobiographical memories (Noreen et al., 2016; Noreen & Macleod, 2013; Stephens et al., 2013). However, most of this research focusses on the after-effects of suppression, showing enhanced subsequent forgetting of suppressed memories even when those memories contain emotional and self-relevant content. Evidence on how emotion affects the intrusiveness of memories during suppression attempts is inconclusive (Davidson et al., 2020; van Schie & Anderson, 2017) and there is, to our knowledge, no previous research measuring intrusiveness of self-relevant and complex autobiographical memories in this paradigm. Furthermore, although some studies have investigated the fMRI correlates of self-relevant materials (Benoit et al., 2016; Noreen et al., 2016), ours is the first EEG investigation of whether emotional autobiographical memories can be suppressed. This approach is important to understand how we engage memory control in everyday life.

In this experiment, we investigated whether and how people can supress autobiographical memories of their own morally wrong actions. Associated with guilt and shame, such memories threaten people’s view of themselves as morally righteous, and should be particularly relevant targets for motivated memory control (Anderson & Hanslmayr, 2014; Stanley & De Brigard, 2019). Indeed, some findings suggest that memories of one’s own immoral actions become obfuscated over time (Kappes & Crockett, 2016; Kouchaki & Gino, 2016), as such memories are more likely to be forgotten over a delay, and if remembered, are rated as less vivid and more distant in time (Escobedo & Adolphs, 2010) than memories of moral actions. However, more recent investigations were unable to replicate some of these findings and these researchers (Stanley et al., 2018) argued that one’s own immoral actions are likely to be more, rather than less memorable than other types of events. We propose here that these seemingly contradictory ideas are compatible with theoretical accounts of motivated forgetting (Anderson & Hanslmayr, 2014; Stanley & De Brigard, 2019). This theory posits that people are highly motivated to suppress retrieval of intrusive, negative memories to maintain good mental health, and that intrusive memories are likely to recruit inhibitory control to keep those memories out of mind. Such efforts should lead to increased suppression-induced forgetting (Anderson & Hanslmayr, 2014; Benoit et al., 2015; Levy & Anderson, 2012). Therefore, if memories for immoral actions are highly memorable and intrusive (Stanley et al., 2018), then this may indeed make them likely targets of motivated forgetting over the longer term (Kouchaki & Gino, 2016).

To test these hypotheses, we adapted the Think/No-Think paradigm to study autobiographical memories, inspired by previous studies (Noreen & Macleod, 2013; Noreen et al., 2016; Stephens et al., 2013). Participants first remembered and described several autobiographical memories involving morally wrong and morally right actions that they committed. They created a title for each memory, to be used in a later Think/No-Think task as a reminder of the memory. The Think/No-Think task was conducted 24 hours later. In this task participants performed trials in which they received a reminder to one of their autobiographical memories. When a given reminder appeared in green, participants were asked to retrieve the associated event and keep it in awareness for the duration of the trial; but when a reminder appeared in red, they were asked to focus their attention on the reminder, but to exclude the unwelcome memory from awareness, as in the typical paradigm (Anderson et al., 2004; Anderson & Green, 2001). The reminder to a given morally right or morally wrong memory was repeated multiple times throughout the Think/No-Think phase and, across repetitions, was consistently suppressed or retrieved. Critically, we measured memory intrusions on a trial-by-trial basis by asking participants to report, immediately after the trial had ended, whether the associated memory had entered awareness at all during the preceding trial (Benoit et al., 2015; Hellerstedt et al., 2016; Levy & Anderson, 2012).^2^

We measured ERPs and oscillatory EEG power in the theta, alpha and beta bands during the Think/No-Think task to delineate the neurocognitive mechanisms associated with autobiographical retrieval and suppression. We sought to investigate how those neural markers might be modulated by the moral nature of the memories and by intrusions. Rich autobiographical memories are likely to be retrieved over a more protracted time-frame and may involve partially different/additional cognitive processes (see e.g., Sheldon et al., 2019) compared to remembering simple item associations encoded in the same lab session. Of the few studies that have investigated the neural mechanisms of autobiographical memory retrieval with EEG, most have found that autobiographical retrieval is associated with later EEG effects, including late parietal positive ERPs that may reflect conscious recollection (Renoult et al., 2015) but also other late slow drift ERP effects (e.g., Conway et al., 2001; see also discussion in Staresina & Wimber, 2019). No prior research has however described the EEG correlates of autobiographical memory suppression and intrusions, and it was unclear whether such EEG effects would have the same topography and timing as suppression and intrusion effects found for simpler stimuli. We therefore used a data driven approach involving cluster-based permutation tests (Maris & Oostenveld, 2007) to assess differences in ERPs and EEG oscillations between conditions across the whole scalp and time-period after Think and No-Think reminders were shown.

Morally wrong memories have been theorised to be memorable and vivid (Stanley et al., 2018; Stanley & De Brigard, 2019). Therefore, if intrusiveness of a memory is related to its vividness, such memories could trigger more intrusions than morally right memories during suppression attempts. Based on evidence that autobiographical retrieval involves at least partially overlapping neurocognitive processes with simpler forms of episodic recollection (Hebscher et al., 2020; Renoult et al., 2015; Tanguay et al., 2018) we predicted that suppression would reduce recollection-related EEG activity, which would be manifest as a reduced late parietal ERP positivity (Bergström et al., 2007) and reduced oscillatory power in the theta band (Crespo-García et al., 2021; Waldhauser et al., 2015) for No-Think compared to Think reminders. When autobiographical memories intrude into participants’ awareness despite suppression attempts, such trials may be associated with increased parietal ERP positivities and theta-band oscillatory power, if autobiographical memory intrusions recruit recollection-related neurocognitive processes. If morally wrong memories intrude more often than morally right memories, Think vs. No-Think differences in recollection-related activity could therefore be smaller for the former. However, since Hellerstedt et al., (2016) did not find enhanced parietal ERP positivities when simple word-pair memories intruded into awareness, the retrieval processes that are engaged during memory intrusions may be qualitatively different from those engaged during voluntary episodic recollection. Therefore, it was not clear how EEG during intrusion trials would compare to retrieval-related activity during Think trials.

Because the inhibitory control mechanism involved in retrieval suppression is considered domain general (Anderson & Floresco, 2021; Anderson & Hulbert, 2021; Apšvalka et al., 2022; Depue et al., 2016; Gagnepain et al., 2014, 2017; Hu et al., 2017), EEG correlates of autobiographical memory control may be similar to those involved in suppressing simpler memories. Therefore, EEG correlates of autobiographical memory control may be reflected in early ERP negativities (Bergström et al., 2009b; Crespo-García et al., 2021; Mecklinger et al., 2009; Streb et al., 2016) and later sustained alpha/beta power reductions as in prior research (Legrand et al., 2020; Lin et al., 2021; Quaedflieg et al., 2020; Waldhauser et al., 2015). Theoretically, recollecting an autobiographical memory involves a slow, gradually unfolding retrieval process (Hebscher et al., 2020; Staresina & Wimber, 2019). Given this, EEG effects associated with both recollection and control processes were predicted to emerge later and, be more sustained than effects found in prior research investigating suppression of recently learned item associations.

## Method

### Participants

Thirty-four students between the ages 18 to 21 years (24 females; *M_age_* = 19.15; *SD_age_* = 0.78) at the University of Kent completed the study in exchange for a combination of course credits and *£*5. This sample size was chosen to be either similar to or larger than samples used in the most relevant previous EEG research (e.g., *N* = 32 in Hellerstedt et al., 2016; *N* = 24 in Waldhauser et al., 2015) and provided >80% power to detect a medium effect size of Cohen’s *d*=0.5 (EEG effect sizes associated with suppression are typically in the range of medium to large). All participants self-reported that they were right-handed, had normal or corrected-to-normal vision, were psychologically and neurologically healthy and were not taking any psycho-active medication. They were also advised to not take part if they were feeling low or depressed due to the upsetting nature of the memories. Participants provided informed consent before taking part. The study was approved by the University of Kent’s School of Psychology ethics committee.

### Design Materials and Procedure

We ran this study in two sessions. In the first session, participants generated autobiographical memories and associated titles for each memory. We used these titles as retrieval cues in the TNT task in the second session, conducted 24 hours later (see Figure 1).

**Figure 1.**
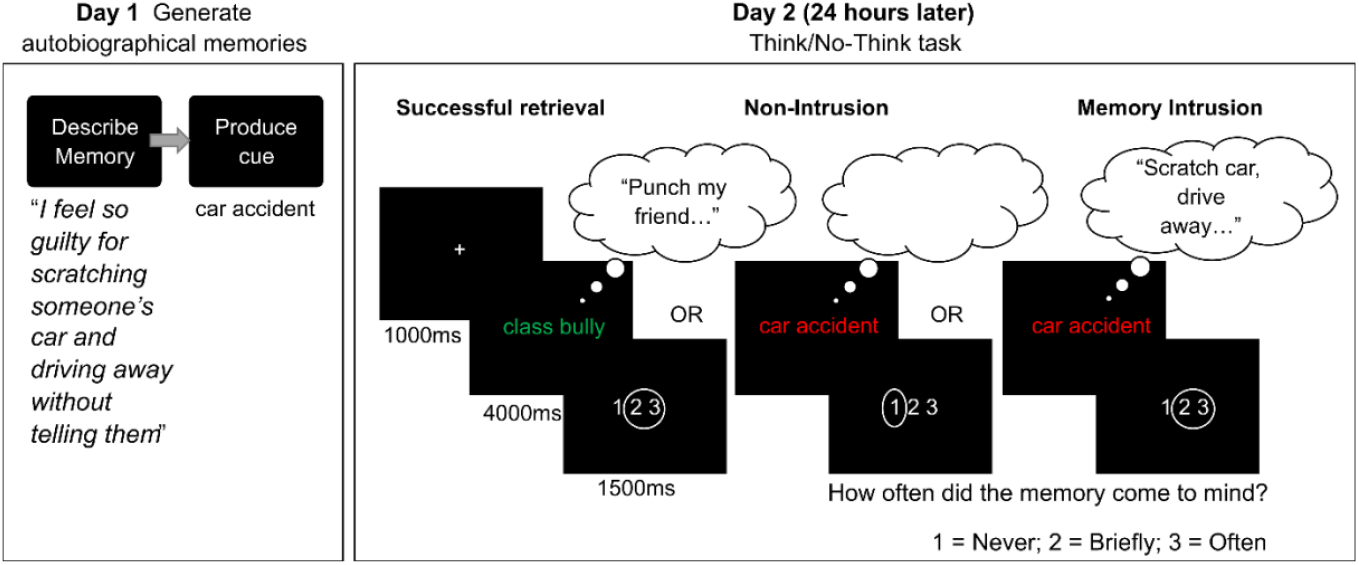
Illustration of the procedure and trial structure of key phases in the experiment. Participants attempted to suppress (No-Think) memories associated to red cues, but consciously recollect the associated memories in response to green cues. Participants indicated how often the associated memory came to mind (retrieval) using the “1 2 3” rating scale.

#### Session one

The first author conducted this session in a lab using the online Qualtrics survey software. We instructed participants to think of 22 different autobiographical memories, one at a time (10 morally wrong and 10 morally right, plus two filler memories involving one birthday and one holiday) and to type a description in a text box provided to them on a computer screen. To aid recollection of morally wrong actions, we instructed them to think of memories in which they lied or cheated, physically or emotionally harmed someone, or in which they committed any other act they considered morally wrong. We also provided similar examples for remembering morally right actions, such as memories in which they were truthful, in which they helped someone physically or emotionally, or in which they committed other morally right actions. Participants had three minutes to think of and write about a specific memory in two to three sentences by describing their actions, the persons involved, the location, and how they felt. We instructed them to avoid writing about events that easily blend with other memories and to describe each event in as much detail as possible.

Next, we instructed participants to think of a unique and specific personal title (henceforth referred to as the “cue”) that would help them to recollect the same memory in the next session. We also advised participants to avoid titles that could evoke multiple memories.

After participants generated the cue, they also rated the memory’s age and vividness, the intentionality and morality of their actions, and their emotional response to remembering the event. These methods and results are reported in a supplementary file (see supplementary file, to be uploaded on OSF after publishing).

#### Session two

The second session began with control and practice tasks to ensure the validity of the methods. We conducted an initial cue-test phase to ensure that participants could remember the autobiographical memory associated to the cue. They self-reported how well they could remember the associated memory in response to a cue presented on the screen. Any memories that participants failed to remember were excluded from further analysis (on average 2.2 memories were excluded in total). Next, they extensively practiced the Think/No-Think task and intrusion ratings over three separate stages using filler memories, until they understood and were following all instructions (see supplemental materials for details).

Think/No-Think phase. In the next phase, the 10 morally wrong and 10 morally right cues were pseudo-randomly assigned to the Think and No-Think conditions in equal proportions by the software, resulting in five cues in each of the four conditions (Morally Right Think; Morally Right No-Think; Morally Wrong Think; and Morally Wrong No-Think). EEG was recorded during this phase.

On each trial, a white fixation cross appeared on a black background for 1000ms, followed by a green or red cue for 4000ms. We instructed participants to think of the associated memory for the entire time that the cue was on the screen, whereas if it appeared in red (No-Think) they were instructed to prevent any thoughts of the associated memory from coming to mind. A direct suppression strategy was used to achieve suppression, which involved two requirements: (1) if the memory happened to intrude, participants were asked to push it out of awareness, while paying full attention to the cue the whole time it was on the screen and (2) participants were asked to refrain from generating substitute memories, images, or words to distract themselves (Benoit & Anderson, 2012; Bergström et al., 2009b).

A 200ms black screen followed each cue, before an intrusion rating scale appeared for 1500ms. The scale presented the options “1 2 3”, and participants selected their option using a keyboard button press. For both Think and No-Think trials, participants were asked to rate a 1 if the associated item never entered awareness during the trial; a 3 if it was in awareness for the entire trial or repeatedly; and a 2 if they only thought the associate briefly. Note that during Think trials, participants ideally would select 3, given their instructions to think of the item and retain it; and on No-Think trials, they would ideally select a 1, given their instructions to suppress the item. An intrusion occurred on No-Think trials if the participant selected a 2 or 3, despite efforts to suppress retrieval.

Cues appeared in a pseudo-random order, one at a time, ensuring that cues from the same Think/No-Think condition were not presented more than three times in a row, in line with previous research (Hellerstedt et al., 2016). After all 20 cues had been presented, a new pseudo-randomised order was generated and all cues were presented one at a time again, and this process was repeated 16 times (leading to a total of 320 trials). Participants were given a short break after each set of 40 trials.

#### Surprise memory test phase

Participants also completed a surprise final recall test where they again described all autobiographical memories after the Think/No-Think phase had been completed using Qualtrics software. In addition, they rated the memories again on the same characteristics that they rated them on in the first session. Final recall and self-reports were analysed but showed no significant effect^3^ of the Think/No-Think manipulation on memories. These methods and results are therefore presented in a supplementary file (to be uploaded on OSF after publishing).

At the end of the second session, participants also completed a compliance questionnaire (adapted from questionnaires typically used in earlier studies such as Hu et al., (2015) to ensure they had completed the task as instructed (see Liu et al., 2021). All participants were compliant with the instructions and were therefore included in further analyses.

### EEG recording and pre-processing

The EEG data were recorded with a sampling rate of 500Hz and bandpass of 0.05-70Hz using 30 Ag/AgCl electrodes fitted in an EasyCAP, amplified with a BrainAmp DC amplifier. Data were recorded with an average reference, and AFz was used as the ground electrode. Electrodes were placed below and above the right eye to measure vertical eye movements, and on both left and right outer canthi to measure horizontal eye movements. Impedances were reduced below 5 kΩ before starting the experiment by gentle abrasion of the scalp using cotton buds and saline gel.

EEG data were pre-processed and analysed using the EEGLAB toolbox (Delorme & Makeig, 2004) for Matlab version R2018b and self-written code. For each participant, data from all electrodes were first re-referenced offline to the average readings of the left and right mastoid electrodes. A 0.1Hz high-pass filter was applied to the re-referenced data. Then, the continuous EEG data was divided into epochs beginning 1000ms pre-stimulus-onset and ending 4000ms post-stimulus onset, baseline corrected on the −200-0ms prestimulus period. This data was visually inspected to delete epochs with extremely noisy data or channels. Independent component analysis (ICA) was then conducted to identify and remove ocular and muscle related components. Then, a 30Hz low-pass filter was applied to the data. Finally, any remaining epochs with large noise were removed based on visual inspection, and epochs were baseline corrected again against the −200-0ms pre-stimulus period for ERP analysis.

### EEG data extraction

We derived two complementary measures from the EEG epochs. First, we computed ERPs by averaging over all trials in each condition for each participant. Second, the single trial epochs were submitted to a time-frequency decomposition in FieldTrip (Oostenveld et al., 2011), using a complex Morlet wavelet transform to decompose the EEG into estimates of oscillation power across different frequencies and timepoints. Twenty-seven wavelets with centre frequencies ranging between 4-30Hz (in steps of 1 Hz), were convolved with the EEG data to produce power estimates across time-steps of 5ms. Each wavelet had a width of three cycles to prioritise temporal resolution over frequency resolution. To remove edge artefacts, epochs were truncated to −625 to 3500ms. A pre-stimulus baseline period of −625 to −325ms was used to normalise the oscillatory power to decibels (dB), since using a baseline period closer to stimulus presentation may lead to artificial “bleeding” of post-stimulus activity into the baseline period.

### EEG statistical analysis

The timing and scalp distribution of EEG effects associated with autobiographical retrieval suppression is largely unknown (Hebscher et al., 2020), so, statistical analyses investigated possible condition differences across locations and time without focusing on pre-determined time-points or electrodes of interest, while also applying a threshold to control for false positives^4^. Data from all 28 scalp electrodes and all post-stimulus time-points were split in two separate time-windows (0ms to 1750ms and 1750ms to 3500ms) which were submitted to nonparametric cluster-based permutation tests using the FieldTrip toolbox (Oostenveld et al., 2011). This approach was used for both ERP amplitudes and oscillatory power derived from time-frequency decomposition, wherein frequency (ranging from 4-30Hz) was added as a third dimension along with electrode locations and time-points.

In the first stage, key experimental conditions (see below) were compared by conducting t-tests at every time (and frequency) point at each electrode site, and adjacent data points in the 2-D (ERP) or 3-D (Oscillation) space that showed significant differences at an uncorrected threshold (*p* < .05) between conditions were identified and grouped into clusters. A minimum of two neighbouring electrodes needed to have significant t-test results to be considered a cluster (default parameter). A cluster-statistic was calculated by summing all the t-values in each cluster. In the second stage, a permutation test was applied to the data to determine which of the observed clusters were statistically significant at a corrected threshold. This test involved randomly reshuffling the condition labels using the Monte-Carlo resampling method (5000 permutations) and calculating a null distribution of cluster-statistics. The observed cluster-statistics were compared against the null distribution to calculate a *p*-value for each cluster. The distribution of significant clusters in time, space and frequency dimensions was then used to determine when, where and in which EEG frequencies the conditions differed. However, it is important to note that the precise edges and peaks of effects in the spatiotemporal and frequency dimensions cannot be determined with this method (see Maris & Oostenveld, 2007; Hellerstedt et al., 2021; Sassenhagen & Draschkow, 2019) for a detailed explanation of the cluster-based permutation test analysis.

For both ERPs and oscillations, the main analysis was conducted to test for differences in neural activity when retrieving versus suppressing autobiographical memories, and to investigate whether such Think/No-Think neural effects differed for morally wrong versus right autobiographical memories. For this analysis, we divided epochs into separate conditions based on a 2 (Instruction Type) x 2 (Memory Type) design in each half of the Think/No-Think task (i.e., the first 8 repetitions of cues corresponded to the first half of trials, and the subsequent 8 repetitions of cues corresponded to the second half of trials) (see (Depue et al., 2013; Hanslmayr et al., 2009; Hellerstedt et al., 2016). On average, 36 trials remained in each condition after pre-processing the data (with individuals contributing between 17-40 trials). We applied this main 2×2 factorial analysis separately in the first half and second half of Think/No-Think trials to investigate if ERP and EEG oscillation effects related to retrieval suppression and the moral nature of memories were present or absent in the early versus late parts of the task. If the interaction between factors was significant, we tested the effects of the Think/No-Think manipulation separately for each memory type.

We also conducted a complementary analysis of ERPs and EEG oscillations to investigate the neural correlates of autobiographical memory intrusions. We categorised epochs into three conditions based on Think vs. No-Think condition and participants’ introspective reports of retrieval using the intrusion rating scale. The conditions were: Successful Retrieval, if participants indicated that memories came to mind briefly or often in the Think condition; Intrusions, if participants indicated the memories came to mind briefly or often in the No-Think condition; and Non-Intrusions, if participants reported that the memories did not come to mind in the No-Think condition. This analysis was collapsed across morally right and wrong memory types and TNT halves to ensure sufficient trial numbers for adequate signal-to-noise ratio. Participants with low trial numbers in the intrusions condition were excluded. Thus, only 26 (out of 34) participants qualified for the trial number cut-off (more than 13 intrusion trials) in this analysis. There were on average 130 (range 79-158) trials in the successful retrieval condition, 101 trials on average (range 69-131) in the Non-Intrusions condition, and 31 trials on average in the intrusions condition (range 13-64) after pre-processing the data. We conducted three pairwise comparisons for both ERPs and oscillations: i) Successful Retrieval vs. Non-Intrusions; ii) Intrusions vs. Successful Retrieval, and iii) Intrusions vs. Non-Intrusions.

## Results

### Behavioural Results

#### Memory retrieval and intrusions

Participants’ trial-by-trial ratings during the Think/No-Think task were used to calculate the frequency of trials accompanied by awareness of the associated memory during Think trials and No-Think trials, separately for Morally right and Morally wrong memories within both the first half of the TNT phase and the second half. Specifically, we computed the percentage of trials where retrieval occurred; because ratings of “briefly” or “often” involved retrieval, we combined trials with those ratings together, following prior work with this method (Hellerstedt et al., 2016; Levy & Anderson, 2012).

Given the differing instructions across the Think and No-Think conditions, we take self-reported retrieval judgments in the Think condition to indicate voluntary retrieval, whereas we interpret self-reported retrieval in the No-Think condition as intrusions. Because these conditions may measure distinct cognitive processes, a 2 (Memory Type) x 2 (TNT half) repeated measures ANOVA was conducted separately for the ratings in the Think vs. No-Think conditions. We used these percentages to test (a) overall differences in intrusion frequency across morally right and morally wrong memories, (b) whether repeated suppression trials reduced intrusion frequency, as is typically found, and any interactions of these factors. See Fig. 2 for an illustration of these results.

**Figure 2.**
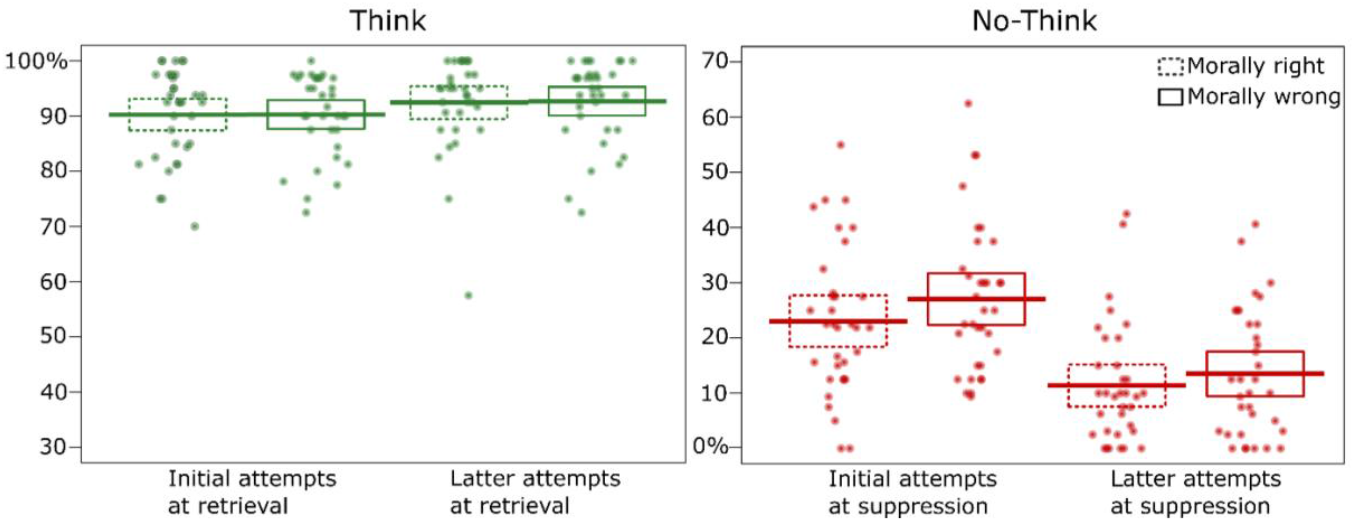
Percentage of trials eliciting memory retrieval compared across memory type and halves of TNT task (first half/initial attempts and second half/latter attempts) in the a) Think condition, reflecting retrieval success; and b) No-Think condition, reflecting intrusion frequency. The dots show the percentage of trials with retrieval for each individual. The thick lines show the group means and the boxes depict the 95% confidence interval of the group means.

Retrieval success increased over repeated retrieval attempts. Retrieval reports indicated that recall success was very high overall, but nevertheless improved between the first half (*M* = 90.26%, *SD* = 6.86%) of the Think/No-Think phase to the second half (*M* = 92.56%, *SD* = 7.20%), *F*(1,33) = 4.52, *p* = .041, partial η^2^ = .12 (Fig. 2a). No overall differences in retrieval success arose for morally right and morally wrong memories, *F*(1,33) = .03, *p* = .87, partial η^2^ = .001), nor was there any interaction between memory type and TNT half, *F*(1,33) = .02, *p* = .89, partial η^2^ = .001). Thus, intentional retrieval was similarly successful for morally right and wrong memories.

Repeated suppression attempts reduced intrusions and morally wrong memories were more intrusive than morally right memories. Morally wrong memories intruded more frequently (*M* = 20.28%, *SD* = 11.43%) than morally right memories (*M* = 17.22%, *SD* = 11.47%), as indicated by the main effect of memory type: *F*(1,33) = 4.62, *p* = .039, partial η^2^ = .123. More memory intrusions occurred during the first (*M* = 25.06%, *SD* = 12.48%) than the second half (*M* = 12.44%, *SD* = 10.34%) of the TNT task, as reflected in the main effect of TNT half: *F*( 1,33) = 78.51, *p*<.001, partial η^2^ = .704. There was no significant interaction between the memory type and TNT half *F*(1,33) = 1.56, *p* = .22, partial η^2^ = .045.

### ERP results

Figure 3 illustrates the grand average ERPs for the Think/No-Think conditions in the first and second half of trials, separated by morality of the memories, together with topographic maps depicting ERP amplitude differences and *t*-values for key factorial and pairwise comparisons.

**Figure 3.**
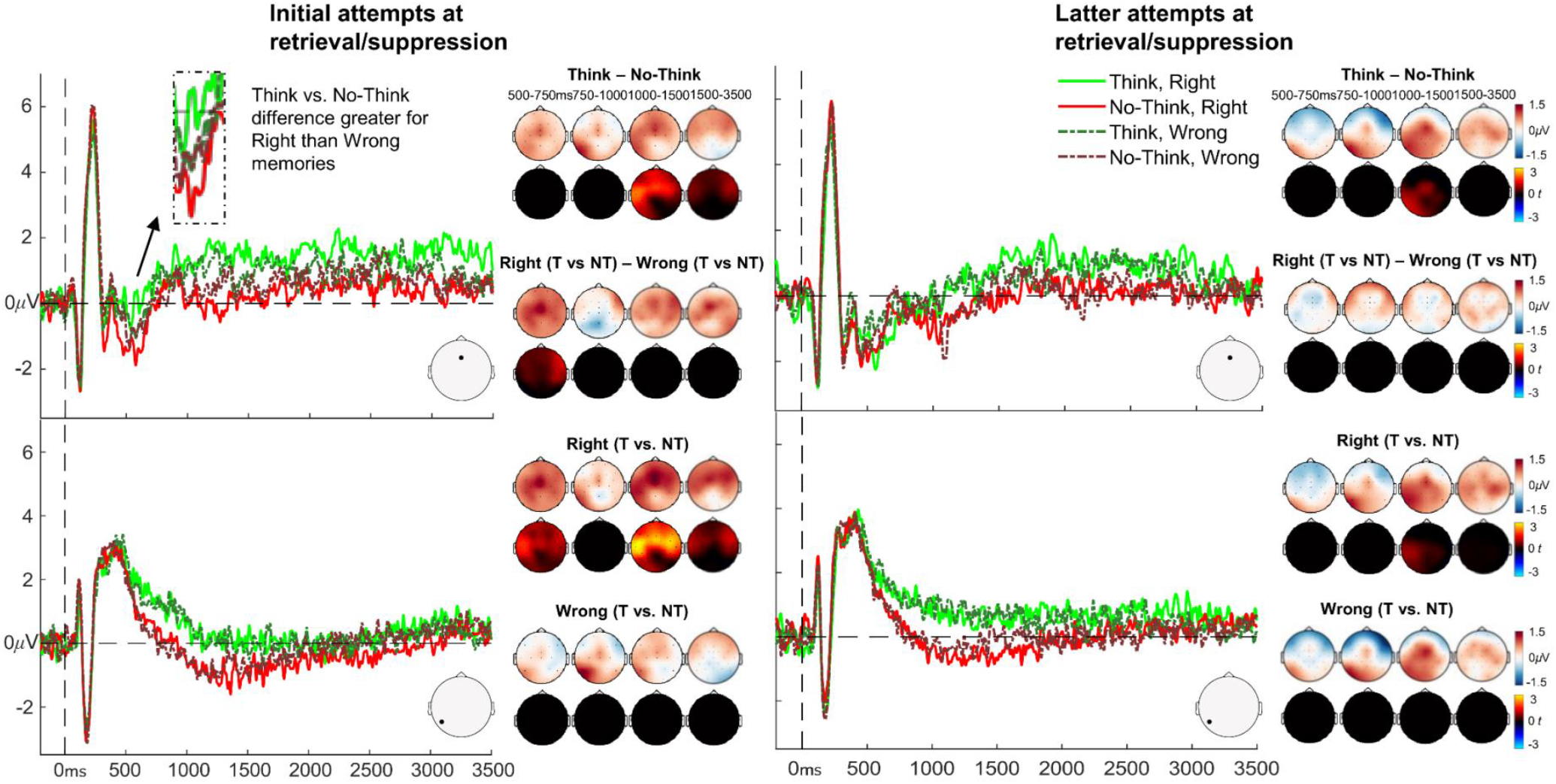
Illustration of grand average ERPs and results from the cluster permutation tests. The ERP graphs show the experimental conditions at mid frontal (Fz, top) and left parietal (P7, bottom) electrode sites during first half (initial attempts) and second half (latter attempts) of the TNT task. Topographic maps on the top rows illustrate mean ERP amplitude differences between conditions (blue/white/red colourmap, in μV) and topographic maps on the bottom rows illustrates mean *t*-values (blue/black/red colourmap, in *t*) for the condition differences as generated through the cluster-based permutation tests. The colour scale represents magnitude and direction of the effect. *T*-maps have been thresholded to only show significant clusters.

#### First half ERPs related to retrieval and suppression of morally wrong and right memories

Overall, we did not find significant clusters for the morally right vs. wrong memory type comparison, but importantly, there were significant clusters showing differences in ERPs between Think vs. No-Think conditions and the interaction between Instruction Type and Memory Type revealed significant clusters, as described below.

##### Overall, suppression attenuates retrieval-related ERPs

We found four significant positive clusters from 1080 to 1360ms (*p* <.001), 1500 to 1670ms (*p* = .005), 2990 to 3160ms (*p* = .005), and 3210 to 3390ms (*p* = .015), indicating more positive sustained ERP amplitudes during retrieval compared to suppression. These effects were present across left parietal (in the 1080-1360ms cluster), frontal, and central regions (in all clusters) as can be seen from the *t*-value distributions in Figure 3. Thus, consistent with our predictions, suppression attempts reduced parietal ERP positivities, and such effects were later in time and more prolonged than previously found with simpler stimuli.

##### Think vs. No-Think ERP differences are present for morally right but not morally wrong memories

The Think/No-Think x memory type interaction analysis showed that the Think vs. No-Think ERP differences was significantly greater for morally right than morally wrong memories between 550-700ms (*p* = .023). Follow up analyses confirmed that this interaction was driven by a Think > No-Think ERP effect for morally right memories around the same time as the interaction cluster (550-700ms, *p* = .009), whereas no significant clusters were found for morally wrong memories.

Similar positive Think vs. No-Think clusters were also present for morally right memories from around 1080 to 1580ms (*p* < .001), 2150 to 2340ms (*p* = .017), and 2940ms to 3400 (*p* < .001) in frontal and central regions, whereas there were no such significant effects for morally wrong memories (see Fig. 3), indicating that the overall Think vs. No-Think differences were primarily driven by effects for morally right memories.

#### Second half ERPs related to retrieval and suppression of morally wrong and right memories

In the second half of trials, we only found only one significant cluster, revealing more positive ERPs for Think compared to No-Think items, from around 1000-1230ms (*p* = .013) across left-posterior and central regions (Fig. 3). This analysis therefore showed that suppression of autobiographical memories reduced late left parietal positivities also in the second half of trials, in line with our predictions. There were however no significant ERP differences between right vs. wrong memory types, nor any significant clusters for the memory type x Think/No-Think instruction type interaction, suggesting that the neurocognitive processes engaged during retrieval and suppression did not reliably differ for morally right and wrong memories in the second half.

We also tested if the Think vs. No-Think positivity between morally right and morally wrong memories found in the first half was significantly greater than the second half, but the cluster did not reach the threshold for significance (*p* = .04, which is non-significant at the two-tailed alpha =.025 threshold).

#### Intrusion-related ERPs

Successful retrieval elicited more positive ERPs than non-intrusions from around 950ms, lasting until the end of the epoch (3500ms, see Fig. 4). This was reflected by significant positive clusters from 950 to 1340ms (*p* = .001), 1350 to 1580ms (*p* = .001), and 1750 to 3500ms (*p* < .001). This effect was spread across left posterior regions in the early clusters, and in the central and anterior regions across all clusters.

**Figure 4.**
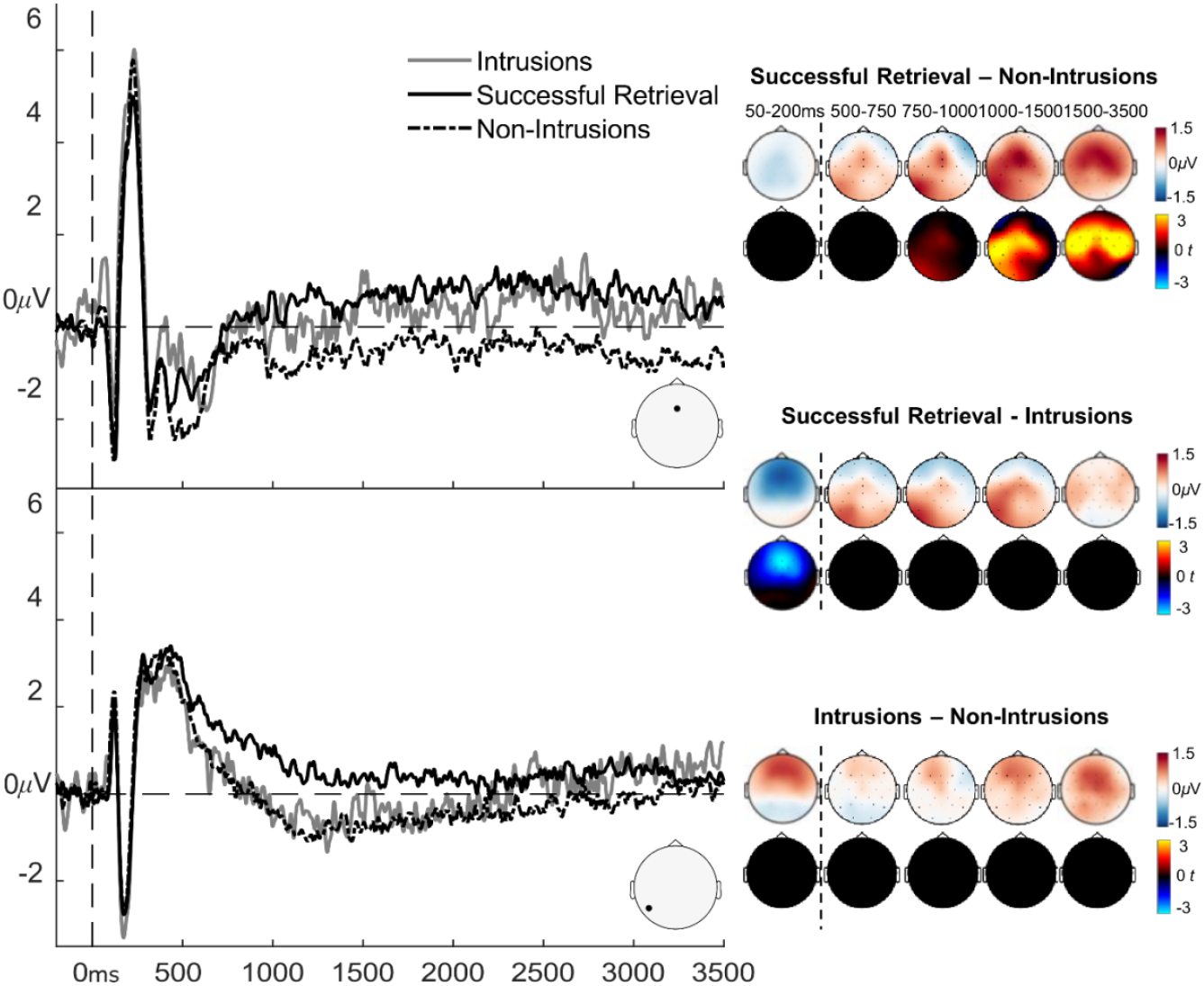
Illustration of grand average ERPs related to autobiographical memory intrusions, and results from the cluster permutation tests comparing the intrusion ERP conditions. The ERP graphs show the experimental conditions at mid frontal (Fz, top) and left parietal (P7, bottom) electrode sites. Topographic maps in the top row show the mean ERP amplitude differences between conditions (blue/white/red colourmap, in μV) and topographic maps in the bottom row illustrates t-values (blue/black/red colourmap, in t) for the differences. The colour scale represents magnitude and direction of the effect. T-maps have been thresholded to only show significant clusters.

When comparing successful retrieval vs. intrusions, one significant negative cluster arose from 50 to 200ms (*p* = .01), caused by more positive ERPs for intrusions than retrieval trials across anterior regions. There were no significant clusters when comparing intrusions and non-intrusions however. The lack of significant differences between intrusions and the other ERP conditions may be explained by the low number of trials for intrusions compared to non-intrusions and successful retrieval trials (see Methods section), leading to noisy intrusion-related ERPs, which may have resulted in low statistical power.

### EEG oscillation results

Figure 5 reports time-frequency plots from a right parietal (P4) electrode for the Think/No-Think conditions in the first and second halves of the TNT phase, separated by morality of the memories, together with topographic maps depicting oscillatory power (dB) differences and t-values for key factorial and pairwise comparisons.

**Figure 5.**
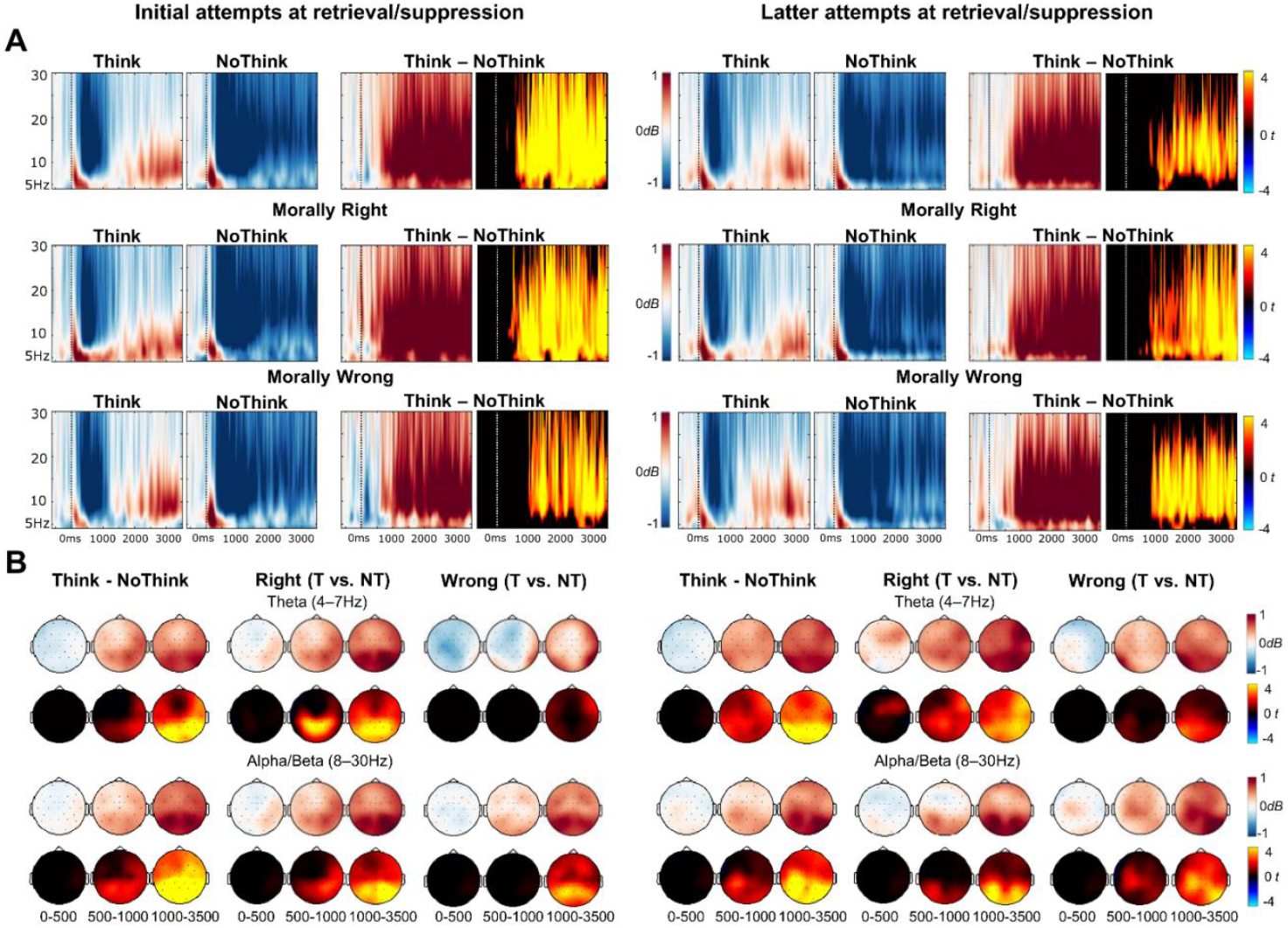
Results of the EEG oscillation analysis comparing retrieval and suppression of morally wrong and right autobiographical memories. (A) Time-Frequency plots from a right parietal (P4) electrode showing the main effect of the Think vs. No-Think manipulation (top row), and the TNT effect separately for morally wrong and right memories (middle and bottom rows respectively) in first half (initial attempts, left panel) and second half (latter attempts, right panel) of the TNT task. Mean power differences are illustrated in the blue/white/red colourmap and t-values for the differences are represented in the cold/black/hot colour map. (B) Topographic maps of power (dB) differences (top rows, blue/white/red colourmap) and t-values for the differences (bottom rows, cold/black/hot colour map) between Think – No-Think conditions at theta (top) and alpha/beta (bottom) frequency bands. T-maps have been thresholded to only show significant clusters.

#### First half EEG oscillations related to retrieval and suppression of morally wrong and right memories

We did not find significant clusters when comparing morally right and wrong memories, either collapsed or when separated between the Think and No-Think conditions. The interaction contrast comparing Think vs. No-Think oscillation differences between right and wrong memories also did not reveal any significant clusters^5^.

We did find a general Think vs. No-Think synchronisation effect across the whole frequency band (4 – 30Hz), with enhanced oscillatory power for Think compared to No-Think trials starting from around 330ms after the reminders were shown, lasting until the end of the epoch 3500ms (*p* < .001). This effect was maximal across parietal scalp regions (see Fig. 5). To assess whether this affect was present for both morally right and wrong memories, we compared Think vs. No-Think trials separately for each memory type. For morally right memories, we found a large Think vs. No-Think synchronisation effect across the whole frequency band (4 – 30 Hz) from 300ms to 3500ms (*ps* < .001), maximal across parietal regions. A similar synchronisation effect was also present for morally wrong memories, but the significant cluster was found later in time, from around 950 to 1750ms and included frequencies from 6 to 30Hz (*p* < .001), whereas from 1750 to 3500ms, the synchronisation effect was significant across the whole frequency range (4 – 30Hz). This effect was also maximal across parietal regions.

#### Second half EEG oscillations related to retrieval and suppression of morally wrong and right memories

As in first half, the interaction contrast comparing Think vs. No-Think oscillation differences between morally right vs. wrong memories was not significant^6^, and there were no main effect differences between morally right vs. wrong memories. A Think vs No-Think synchronisation effect was again found in the second half, which was similar to the effect in the first half (see Fig. 5). The Think vs No-Think main effect cluster showed significantly higher power in the Think than No-Think condition across the whole frequency range (4 – 30Hz) from around 440 to 3500ms post-stimulus (*p* < .001). For morally right memories, the effect was present across the whole frequency band (4 – 30Hz) from around 90ms to 3500ms (*p*s < .001). Similarly, for wrong memories, the significant clusters incorporated the whole frequency band (4 – 30Hz) but the effect started later lasting from 600ms until 3500ms (both *ps* < .001).

Thus, EEG oscillations in both halves were similar and did not vary across memory types, but highly significantly differed depending on whether participants were trying to retrieve or suppress autobiographical memories, with reduced power across theta, alpha, and beta frequencies during suppression compared to retrieval, consistent with predictions.

#### Intrusion-related EEG oscillations

We next compared EEG oscillations across trials based on whether participants successfully retrieved autobiographical memories in the Think condition, successfully suppressed memories in the No-Think condition, or failed at suppression and experienced intrusions of autobiographical memories during No-Think trials. When comparing successful retrieval with non-intrusion suppression trials, a synchronisation effect was present across the whole frequency range (4-30Hz) starting around 400ms to 3500ms (both *ps* <.001), maximal across parietal regions (Fig. 6). Successful retrieval was also associated with a similar synchronisation effect compared to intrusions, again spanning across the whole frequency range (4-30Hz) and starting around 650ms to 3500ms (both *ps* < .001). These differences therefore corresponded closely to the effects of Think/No-Think instruction observed in the earlier factorial analysis of EEG oscillations, with enhanced theta, alpha and beta power during retrieval compared to suppression. Interestingly, when comparing intrusions with non-intrusions, a significant synchronisation effect was found in the alpha and beta bands (7-30Hz), late in the epoch, around 2800ms to 3500ms (*p* = .008). As can be seen in Figure 6, this difference between intrusions and non-intrusions was caused by a more sustained alpha/beta desynchronisation effect during non-intrusion trials. In contrast, during intrusion trials there was an initial alpha/beta desynchronisation effect, but this effect weakened during the latter part of the epoch, where the power in this condition returned to levels similar to the baseline period^7^.

**Figure 6.**
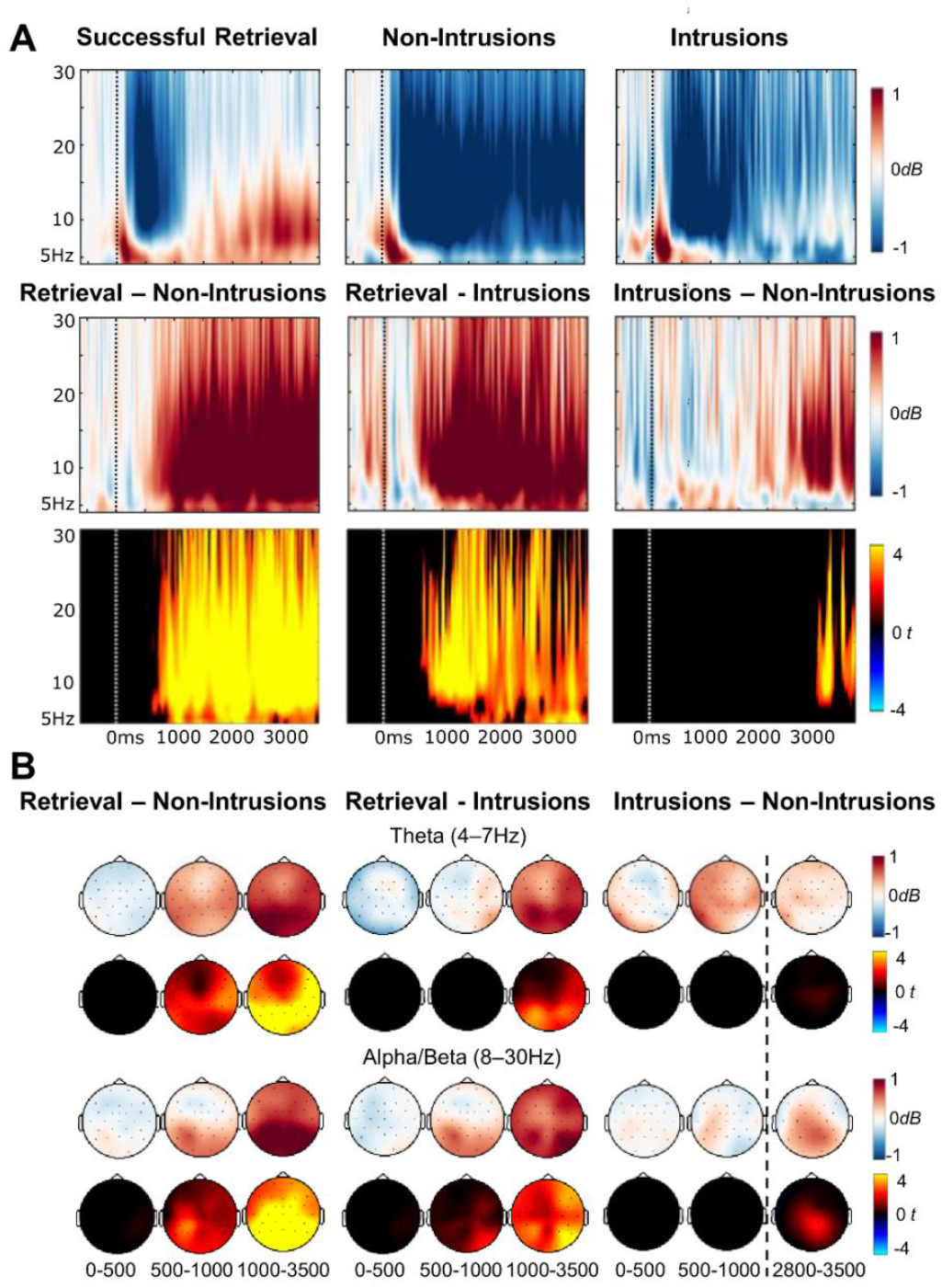
Results of the EEG oscillation analysis comparing intrusions with successful retrieval and successful suppression trials. (A) The top row shows time-frequency plots for each condition separately from a right parietal electrode (P4), and the second and third rows show pairwise differences between conditions. Mean power (in dB) is illustrated in the blue/white/red colourmap and t-values for the differences are represented in the cold/black/hot colour map. (B) Topographic maps below show power (dB) differences (top rows, blue/white/red colourmap) and t-values for the differences (bottom rows, cold/black/hot colour map) between conditions at theta (top) and alpha/beta (bottom) frequency bands. T-maps have been thresholded to only show significant clusters.

## Discussion

In this experiment, we examined the neurocognitive mechanisms underlying the suppression of autobiographical memory retrieval and investigated how the moral nature of memories affects our ability to stop them from intruding into awareness. The results suggest that autobiographical memory retrieval can be suppressed, and that repeated suppression attempts reduce intrusions. Participants’ memories of their morally wrong actions were particularly intrusive compared to their memories of morally right actions, indicating that controlling memories that threaten our self-concept may be especially challenging. ERP and EEG oscillation markers of autobiographical suppression and intrusions were found to be mostly in-line with findings from prior research investigating suppression of simple laboratory materials. However, the time-course of these ERP and EEG effects revealed interesting new evidence of how autobiographical memory retrieval and control processes change over time. Our findings thus provide novel behavioural and neural evidence concerning the control of multifaceted and emotionally charged real life memories.

Behaviourally, we found that the frequency of autobiographical memory intrusions reduced over repeated suppression attempts, a replication of a reliable effect in previous TNT research with simpler stimuli (Benoit et al., 2015; Davidson et al., 2020; Gagnepain et al., 2017; Harrington et al., 2021; Hellerstedt et al., 2016; Levy & Anderson, 2012; Mary et al., 2020; van Schie & Anderson, 2017). Furthermore, morally wrong memories intruded more frequently than morally right memories, in general. Research with the TNT paradigm on how the emotional nature of memories affects their intrusiveness is scarce. In the three studies that have investigated this issue, negative memories were found to be either more intrusive (Davidson et al., 2020), numerically less intrusive (Gagnepain et al., 2017), or not differently intrusive (Harrington et al., 2021) than neutral memories. Participants in this study self-reported that they felt more guilty, ashamed and overall more negative while retrieving morally wrong compared to morally right memories (see supplementary file, to be uploaded on OSF after publishing), suggesting that emotionally negative memories are more intrusive than positive memories. Importantly, this finding provides new insights by investigating this issue with complex autobiographical memories that reflects the types of unwanted memories we often confront in everyday life.

The findings are in line with suggestions that memories of our own morally wrong behaviour may be particularly memorable (Stanley et al., 2018). Intrusive memories are likely to require effortful cognitive control processes to prevent them from coming to mind (Anderson & Hanslmayr, 2014; Benoit et al., 2015), which may make such memories more vunerable to motivated forgetting in the longer term (Kouchaki & Gino, 2015, 2016; Shu & Gino, 2012). However, it’s important to note that we did not observe changes in how the morally wrong (or right) memories were described as a result of the Think/No-Think manipulation in this study (see supplementary file, to be uploaded on OSF after publishing). Most prior evidence for morally motivated *forgetting* comes from testing changes in phenomenology of episodes encoded in the lab, or reduced memory accuracy for imagined events (e.g., Kouchaki & Gino, 2016; Shu et al., 2011) rather than memory for events that people have personally experienced in their everyday life (c.f. Stanley et al., 2018). Some evidence suggests that personal autobiographical memories may not necessarily be completely forgotten after suppression, but may instead change in more subtle ways to become less detailed (Noreen & Macleod, 2013; Stephens et al., 2013). Core details of important events in people’s past may however be more resistant to forgetting, especially those details that they have repeatedly dwelled on over long time periods (Noreen & Macleod, 2013). Our method for assessing memory changes may not have been sufficiently sensitive to such subtle and complex effects. Nevertheless, participants’ intrusion ratings showed that autobiographical memories did become less intrusive over repeated attempts at suppression, suggesting that some changes to those memories may have occurred within the session. Future research should investigate whether these suppression-induced changes to memory intrusiveness are long-lasting and should also use more sensitive measures of autobiographical forgetting.

### ERP correlates of autobiographical memory suppression and intrusions

Successful retrieval suppression led to a sustained reduction in positivity compared to retrieval, that began around 750ms after cue onset and lasted until the end of the epoch at 3500ms. This reduced positivity was initially strongest across left parietal scalp regions but later was maximal across fronto-central regions. Similar but weaker effects arose when contrasting ERPs by Think/No-Think condition regardless of self-reported success, suggesting that these differences were primarily driven by trials where both suppression and retrieval attempts succeeded (around 90-95% of trials for retrieval attempts, and 75-85% of trials for suppression attempts).

The earlier parietal modulation strongly resembles the left-parietal ERP positivity thought to index conscious recollection (Rugg & Curran, 2007; Wilding, 2000) including during autobiographical retrieval (Renoult et al., 2015; Tanguay et al., 2018). A suppression-induced reduction in left-parietal ERP positivity has been widely found in the EEG literature on memory control with simpler stimuli (e.g., Bergström et al., 2007, 2009a, 2009b; Chen et al., 2012; Depue et al., 2013; Hanslmayr et al., 2009; Hellerstedt et al., 2016), and these results therefore replicate this suppression effect with autobiographical memories (see also Bergstrom et al., 2013; Hu et al., 2015). The timing of the parietal ERP reduction was later than is usually found, as it was maximal around 750-1000ms post-stimulus onset, compared to peaks occurring around 500-800ms post-stimulus in previous TNT research with simpler stimuli. This result is consistent with recollection of autobiographical memories unfolding slowly and gradually over time (Conway & Pleydell-Pearce, 2000; Daselaar et al., 2008; Hebscher et al., 2020; Inman et al., 2018; McCormick et al., 2015). Importantly, this finding suggests that suppression can interrupt brain processes that normally contribute to successful autobiographical recollection.

In the later part of the epoch, successful suppression was associated with more negative fronto-central slow waves compared to successful retrieval, and this effect was sustained between around 1500 to 3500ms. Similar frontal slow waves have been found in previous Think/No-Think research with simpler stimuli, and a reduction in frontal positivity during suppression has been suggested to reflect cognitive control processes that are engaged to keep the memory from entering awareness (Depue et al., 2013; Hanslmayr et al., 2009). Therefore, our results provide neurophysiological and behavioural evidence that retrieval of autobiographical memories can be successfully suppressed and may recruit similar neurocognitive processes as the suppression of simpler memories.

In contrast to the large ERP effects related to successful suppression, there was less clear evidence for ERP markers of intrusions^8^. There was a very early difference around 40-200ms across anterior regions with more negative ERP amplitudes for successful retrieval compared to intrusions, and numerically also for non-intrusions vs. intrusions (although the latter comparison was not significant). This time-region is usually reflected by P1 and P2 peaks, that are thought to reflect early selective attention processes (Bergström et al., 2009b; Hellerstedt et al., 2016). Therefore, this effect could indicate *enhanced* attentional allocation towards the No-Think cue on trials when an intrusion occurred, or conversely, *reduced* attentional allocation of attention during non-intrusions, consistent with an explanation whereby early attentional control to cues is important for preventing intrusions.

During the later part of the epoch, whereas non-intrusion ERPs were more negative than retrieval ERPs, ERPs associated with intrusions were not significantly different from either retrieval or non-intrusion ERPs. Numerically, the left parietal ERP effect was in line with Hellerstedt et al.’s (2016) findings. Intrusions elicited similar ERP amplitudes to non-intrusions between ~500-1000ms post-stimulus. However, from ~1000ms onwards across central and frontal scalp regions, intrusion ERP amplitudes fell in between non-intrusions and intentional retrieval, indicating that intrusions made ERPs more similar to intentional retrieval. However, since these effects did not survive statistical thresholding, we do not interpret them further here. The inconclusive statistical results likely occurred due to the overall low number of intrusions, which impaired the signal-to-noise ratio of those ERPs thereby decreasing statistical power for this analysis (which is especially low with conservative cluster-based thresholding). Future research is needed with increased numbers of intrusion trials to detect reliable ERP markers of autobiographical intrusions.

When comparing the effects of suppression on ERPs for morally right vs. wrong memories, we found that during initial attempts (first half of the TNT trials), an ERP positivity during retrieval compared to suppression attempts was found for morally right memories but not morally wrong memories. This Think > No-Think positive difference for morally right memories was most reliable around 500-600ms post-stimulus onset and peaked across central and frontal regions. Although ERP modulations indexing the suppression of recollection often peak over the left parietal scalp (e.g., Bergström et al., 2009a, 2009b), recall-related ERPs can be spread across the whole scalp (e.g., Hellerstedt et al., 2021) and are known to emerge from around 500ms after a reminder is encountered (Staresina & Wimber, 2019). Therefore, the ERP positivity for retrieving compared to suppressing morally right memories likely indexes differences in recollection-related activity for these memories, whereas control over recollection may not have been as successful for morally wrong memories during these initial attempts at suppression. Therefore, in line with behavioural self-reports, ERPs indexed the difficulty in avoiding retrieval of morally wrong but not morally right memories.

During latter attempts at the task (second half of trials), Think/No-Think ERP effects were no longer significantly different between morally right and wrong memories, and instead the results only showed general Think > No-Think ERP positivities, similar to the first half of the trials, as described above. We did not find, however, that this difference between Think and No-Think ERPs across memory types differed reliably across the two halves of the task. Similarly, we did not find significant evidence that the reduction in intrusion rates across halves was larger for morally wrong than for morally right memories, although there was a numerical pattern in that direction. Although these patterns were not statistically significant, previous research has found that cognitive processing during early stages of the TNT task can differ from processing in the latter stages (Depue et al., 2007, 2013; Gagnepain et al., 2017; Hanslmayr et al., 2009; Hellerstedt et al., 2016; Hulbert et al., 2016), and it’s possible that our design was underpowered to detect these changes with our relatively conservative statistical analysis.

### Oscillatory correlates of autobiographical memory suppression and intrusions

The EEG oscillation results further indicated that retrieval of personal autobiographical memories can be suppressed. Overall, we found a strong decrease in oscillatory power during successful suppression compared to successful retrieval across all analysed frequencies (4-30Hz), maximal across parietal regions. This effect began around 500 to 1000ms after the reminder appeared and was sustained for as long as the reminder was presented on the screen (3500ms). This broadband and sustained desynchronisation effect resembles the oscillatory correlates of successful memory suppression described in previous research with simpler episodic memories (Crespo-García et al., 2021; Ketz et al., 2014; Legrand et al., 2020; Lin et al., 2021; Quaedflieg et al., 2020; Waldhauser et al., 2015).

Previous research indicates that memory-related power differences in the theta band (4-7Hz) versus alpha/beta bands (8-30Hz) reflect different cognitive processes. Theta band increases are often related to retrieval success (Hanslmayr et al., 2016; Nyhus & Curran, 2010; Osipova et al., 2006) consistent with our findings of enhanced theta power during retrieval attempts. In contrast, No-Think trials were associated with a significant theta power desynchronisation effect, likely reflecting successful avoidance of retrieval. The results also showed sustained alpha/beta power decreases for No-Think vs. Think reminders. Alpha/beta desynchronization is often found during memory retrieval (see Hanslmayr et al., 2016), and in our results both retrieval and suppression trials were associated with an initial alpha/beta desynchronisation effect within the first second after the cue was shown. Critically however, alpha/beta desynchronization was sustained throughout the epoch – from 1s to the end of the epoch - during memory suppression only, suggesting that the initial versus latter effects were functionally dissociable. Late, long-lasting alpha/beta power decreases for No-Think vs. Think reminders have been argued to reflect sustained control over the memory for as long as the participant is exposed to the reminder (see Waldhauser et al., 2015; Lin et al. 2021). Therefore, the pattern of alpha/beta changes in our study indicates that suppression recruited sustained control processes to prevent autobiographical memories from intruding into awareness.

The theta and alpha/beta reductions described above were found consistently for both morally right and wrong memories, and both during initial and latter attempts at suppression, showing these effects to be highly reliable as an index of autobiographical memory suppression. The lack of EEG oscillation differences between memory types indicates that the oscillations reflected at least partially different neurocognitive processes than the ERPs, which did show differences by memory type. Alternatively, our statistical approach for analysing EEG oscillations may not have been powerful enough to detect subtle differences in neurocognitive processes between morally wrong and right memories that was detectable in the ERP analysis.

Interestingly, correlates of memory intrusions were more apparent in the EEG oscillation analysis, contrasting with the weaker ERP findings for intrusions. Autobiographical memory intrusions were associated with a significant increase in parietal alpha/beta (8-30Hz) power compared to non-intrusion trials, around 2500-3500ms post-stimulus onset. This result contrasts with some previous findings that intrusions *decreased* rather than *increased* oscillatory power compared to successful suppression (Castiglione et al., 2019; Legrand et al., 2020). However, those prior effects were generally observed earlier in time and tended to have more frontal topographies than the intrusion effect found in the present study. Earlier frontal increases in alpha/beta oscillatory power are thought to reflect the engagement of top-down inhibitory processes that are recruited to stop unwanted memories from coming to mind (Castiglione et al., 2019), similar to motor-action stopping.

The much later, more posterior intrusion-related oscillation changes in the current study could reflect a functionally different mechanism. As described above, sustained decreases in alpha/beta oscillatory power has been suggested to index maintenance of control over retrieval (Quaedflieg et al., 2020; Waldhauser et al., 2015). In the current study, both intrusion and non-intrusion trials were associated with an initial decrease in alpha/beta power, but towards the last second of the epoch (around 2.5s after the reminder was shown), alpha/beta power for intrusion trials returned to near baseline levels. In contrast, decreases in alpha/beta oscillatory power were sustained until the end of the non-intrusion trials. Hence, the late intrusion-related increase in alpha/beta oscillatory power could reflect a failure to maintain control throughout the time the reminder was shown (see also Lin et al., 2021). This interpretation is consistent with behavioural findings that intrusions are more likely if the No-Think reminder is shown for longer to participants, suggesting that people’s memory control ability dissipates over time (van Schie & Anderson, 2017). Therefore, the present results provide novel oscillatory evidence that autobiographical memory intrusions could occur due to a lapse in sustained control.

Even though autobiographical memory intrusions clearly did occur as evident by both self-reports and a unique pattern of EEG effects, participants reported experiencing intrusions less frequently in this study (on average 20% of trials) than in previous studies with simpler memories of paired word or picture associates (around 35% on average, e.g., Levy & Anderson 2012). This disparity could be due to relatively weak cue-memory associations in our autobiographical memory version of the paradigm reducing the likelihood that cues would elicit recall, which is a common issue in autobiographical memory research in general (see St Jacques & De Brigard, 2015). In contrast, in memory suppression paradigms using over-learned paired associates, cues may be more likely to reactive the associated memory automatically (see Hellerstedt et al., 2016).

One issue that is not clear from the present study is *why* morally wrong autobiographical memories were more difficult to suppress than were morally right memories. Our findings are in line with predictions from Stanley and De Brigard (2019), who suggested that the guilt, shame, and self-threatening aspects of these memories would make them particularly intrusive. However, further research is required to determine which specific features of morally wrong memories render them more intrusive than morally right memories. Morally wrong memories could be particularly memorable because they threaten our belief that we are inherently good (Stanley and De Brigard, 2019), and are associated with negative emotions of guilt and/or shame. Alternatively, the more intrusive nature of morally wrong memories found in this study could simply be due to the general negative valence of morally wrong memories without being specifically related to morality-relevant memory characteristics. Therefore, further research that separates contributions to intrusiveness from emotions, morality, and how strongly a memory is threatens the belief of a good-self will be imperative to understand why these memories could be more difficult to control in everyday life.

In conclusion, the results of this experiment provide both behavioural and electrophysiological evidence that people can suppress unwanted autobiographical memories as early as half a second after a reminder is encountered. Autobiographical memory suppression can fail however and result in unwanted memories intruding into awareness, which may be caused by a lapse in sustained control over the memory. Such intrusions reduce over repeated attempts however, indicating better memory control with practice and repetition. This pattern of results both converges with and extends previous findings, because it is the first demonstration of control over intrusions of real-life, self-relevant autobiographical memories (but see also Noreen & MacLeod, 2013; Noreen et al., 2016; Stephens et al., 2013). Both behavioural and EEG evidence also suggests that intentional suppression may be more difficult for memories of our morally wrong actions than for memories of our morally right actions. Importantly however, the results indicate that even such personal and emotionally negative memories can be controlled, which may be helpful for mental health and wellbeing in our everyday lives.

## Declarations

### Ethics Approval

Ethics approval was awarded by University of Kent’s, School of Psychology ethics committee.

### Consent to participate & publish

Informed consent was obtained from all individual participants included in the study. They specifically consented for anonymised data to be available on online repositories and to be published in peer-reviewed journals.

### Availability of data, materials, & code

The data, materials, code, and supplementary information are uploaded on the Open Science Framework, an open science platform. The OSF link will be provided after the manuscript is accepted for publication.

### Author contributions

*Conceptualisation* A.S., R.H., Z.M.B.; *Methodology* A.S., R.H., M.C.A., Z.M.B.; *Data collection and investigation* A.S., R.H., Z.M.B; *Formal Analysis* A.S., R.H., Z.M.B.; *Writing – Original Draft* A.S., and Z.M.B.; *Writing – Review & Editing* A.S., R.H., M.C.A., Z.M.B; *Supervision* Z.M.B.

## Acknowledgements

We would like to thank Aderayo Ariyo, Chloe Brunskill, Manahil Aslam, Rachel Arthur, and Samantha Gilbert for their assistance with collecting data for this project. We would also like to thank Brenton M. Wiernik for the pre-print template, which can be found on OSF via this link:https://osf.io/t4eqp/.

## Footnotes

1 Not to be confused with mid-frontal theta power changes that are often thought to reflect cognitive control processes (Waldhauser et al., 2015). Here, we refer to theta activity specifically linked to memory retrieval, which should be absent if retrieval suppression is successful (ter Wal et al., 2021; Waldhauser et al., 2015).

2 Participants also completed a surprise recall test and provided subjective ratings of various features and emotions associated with their memories before and after the Think/No-Think phase to test for changes in memory as a function of the manipulation. However, these analyses did not show suppression-induced changes in phenomenology or descriptions of memories. So, they are presented in a supplementary file for brevity (to be uploaded on OSF after publishing). Further note that baseline items were not used in this study, so a clear test of suppression-induced forgetting was not possible.

3 In brief, final recall performance showed a non-significant but numerical tendency towards enhanced forgetting of suppressed compared to retrieved memories. The lack of significant suppression after-effects could be due to insufficient statistical power/noisy measurement in our design (we only had five memories per experimental condition and recall performance was measured by experimenter ratings of free text memory descriptions).

4 An additional analysis using multifactorial ANOVAs was conducted to compare our results against prior literature (Hellerstedt et al., 2016), uncorrected for multiple comparisons. Overall, these effects were significant across longer time periods and were more broadly distributed across the scalp in the ANOVA results, because of the laxer statistical threshold. The main divergence between cluster-based tests and ANOVAs were found in the intrusion analysis, where ANOVAs showed additional intrusion-related ERP effects that were in line with previous research.

5 When only the theta band (4-7Hz) was included in a focal analysis of the first half, three significant positive clusters were found for the interaction between Think/No-Think instruction and morally right/wrong memories across posterior regions from around 500 to 1000ms (*p* = .01), 1000 to 1300ms (*p* = .01), and 1960 to 2500ms (*p* = .023). The interaction was caused by enhanced theta power for Think vs. No-Think conditions for morally right memories only, whereas there were no significant theta effects for morally wrong memories. Thus, theta oscillations in the first half of the TNT task showed a similar pattern as ERPs when analysed in a less conservative way than was done in the main analysis. The pattern of theta effects was in line with predictions.

6 There were no significant clusters in the instruction type x memory type interaction analysis when only the theta band was included separately in a focal, less conservative analysis in the second block. Thus, like the ERPs, theta oscillations did not differ as a function of memory type in the second half of the TNT task.

7 There were no additional significant clusters when only the theta band was included in a focal analysis of the intrusion conditions.

8 Supplementary ANOVAs however revealed intrusion-related ERP effects, but these results were uncorrected for false-positives so are not reported here (see supplementary file, to be uploaded on OSF after publishing). In brief, the findings were in line with the only previous study on ERP effects related to TNT intrusions by Hellerstedt et al., (2016). A greater FN400 was present for intrusions compared to successful suppression, potentially reflecting reactivation of the memory (see also Bergström et al., 2012; Hellerstedt & Johansson, 2014).

